# *Drosophila* Aop imposes a delay on E(spl)-mediated repression of Ato during R8 specification

**DOI:** 10.1101/678169

**Authors:** Adam T. Majot, Lucas M. Jozwick, Clifton P. Bishop, Ashok P. Bidwai

## Abstract

*Drosophila* retinal patterning requires the expression of Atonal (Ato) through coordinated regulation of 5’ and 3’ enhancer modules. *ato-3’* directs initial expression of Ato which then directs autoregulation via *5’-ato*. Notch (N) signaling also regulates *5’-ato*, first enhancing Ato expression and later repressing Ato by inducing E(spl) bHLHs. N signaling balances these opposing functions by directing its obligate nuclear transcription factor, Suppressor of Hairless (Su(H)), only in repressing *5’-ato*. In this study, we reveal a novel and more nuanced role for Su(H) in its regulation of *5’-ato*. During retinal patterning, Su(H) is required for the expression Anterior open (Aop), which, in turn, promotes *5’-ato* activity. We demonstrate that Aop is induced early in retinal patterning via N pathway activity, wherein Aop is required cell-autonomously for robust Ato expression during photoreceptor specification. In *aop* mutants, expression from both *ato* enhancers is perturbed, suggesting that Aop promotes the Ato autoregulation through maintenance of *ato-3’* activity. Clonal analysis indicates that Aop indirectly opposes E(spl)-mediated repression of Ato. In the absence of both Aop and E(spl), Ato expression is restored and the founding ommatidial photoreceptors, R8s, are specified. These findings suggest that N signaling, through a potentially conserved relationship with Aop, imposes a delay on *ato* repression, thus permitting autoregulation and retinogenesis.

**Author Summary:** The eye of the fruit fly has served as a paradigm to understand tissue patterning. Complex intercellular signaling networks cooperate during retinal development to allow cells to become specialized visual-system precursor neurons at a specific time and place. These neurons are precisely spaced within the developing retina and later recruit other cells to form the repeated units that comprise insect eyes. The exact placement of each precursor cell precipitates from the precise regulation of the *atonal* gene, which is first expressed in a cluster of (10-20) cells before becoming restricted to only one cell from each cluster. The Notch signaling pathway is required for both aspects of *atonal* regulation, first permitting up-regulation within each cluster, and then the subsequent down-regulation to a single cell. However, the connection between these two modes of Notch signaling had remained unclear. In this report, we have identified that the *anterior open* gene is required to impose a delay on the restrictive mode of Notch signaling, permitting the initial up-regulation of atonal to occur freely. In flies mutant for *anterior open*, *atonal* bypasses its own up-regulation and proceeds directly to its singled-out pattern but with significantly diminished robustness than occurs normally.

## Introduction

The *Drosophila* retina is a widely studied genetic model for neuroepithelial organization. Photoreceptor clusters, termed ommatidia, are established during the third larval instar in dorsoventral columns with the passage of the morphogenetic furrow (MF) across the eye-antennal imaginal disc (Wolff and Ready, 1993). Photoreceptors are initially specified in a staggered pattern through the dynamic expression of Ato within the MF (Jarman et al., 1994). Ato is first expressed in a broad dorsoventral (Fig. 1A, stage-1) column, subsequently upregulated within 10-20 cell intermediate groups (IGs, Fig. 1A, stage-2), each of which are later resolved to a single cell, the R8 (stages-3 and -4; Sun et al., 1998, Baker et al. 1996).

**Figure 1.**
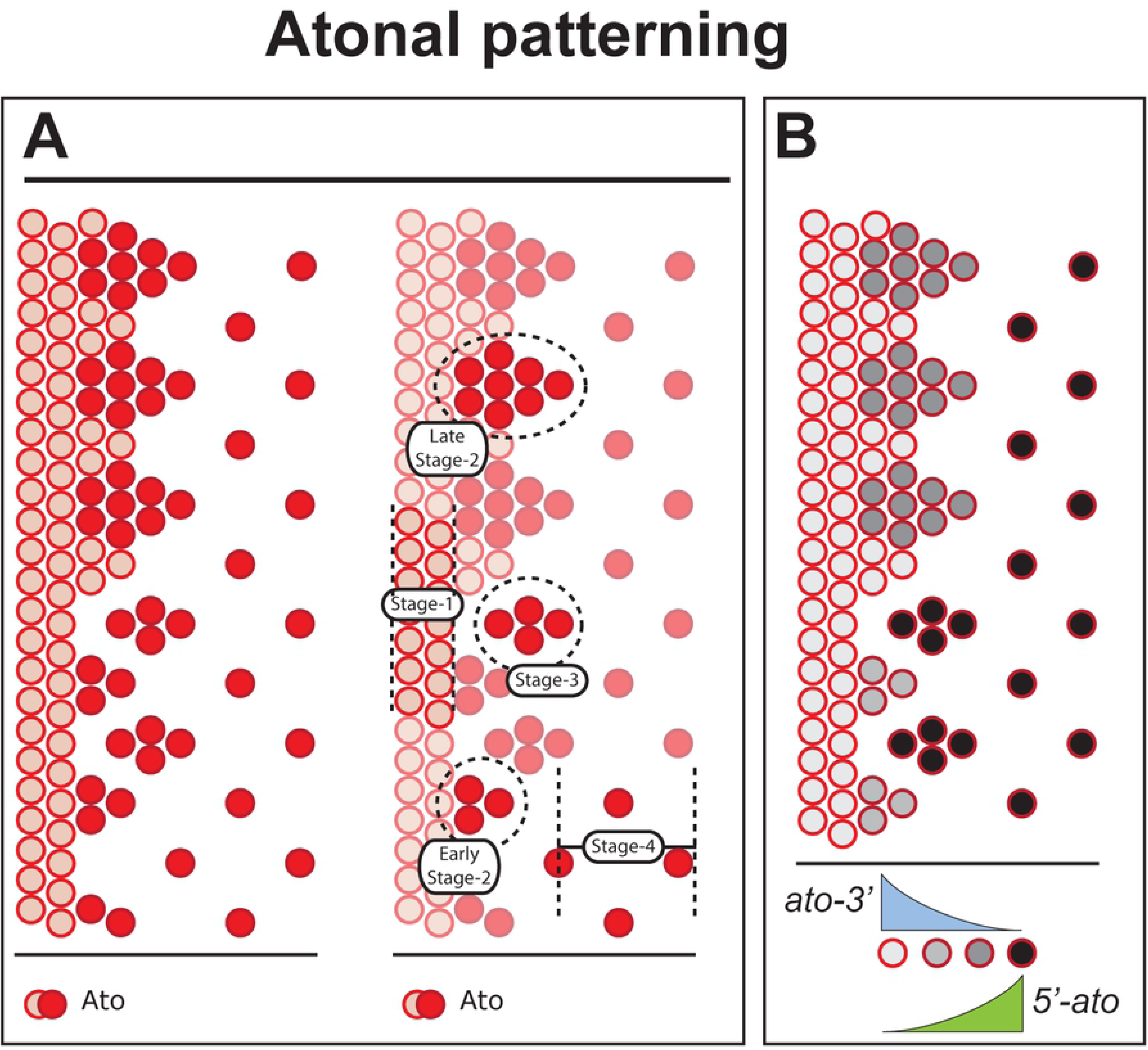
Ato progresses through four distinct stages. (A) Ato is initially expressed in a dorsoventral column of cells via the *ato-3’* enhancer (stage-1). Stage-2 features IG formation (early stage-2) and maturation (late stage-2), with Ato expression undergoing a regulated transition from exclusive dependence on the *ato-3’* enhancer to that of both *ato-3’* and *5’-ato*. Stages 3 and 4 are characterized by the resolution of IGs to individual Ato^+^ R8s, and Ato expression in these final two stages is exclusively dependent on *5’-ato* activity. (B) Ato expression, grey-scaled to illustrate enhancer-specific contributions to Ato expression throughout its expression in the MF.

Ato patterning is dependent upon two enhancer modules, *5’-ato* and *ato-3’* (Fig. 1B), and their complex entanglement with both the N and Epidermal growth factor receptor (EGFR) signaling pathways. Initially, *ato-3’* drives expression of Ato. Ato then induces the N pathway (Baonza and Freeman, 2001), which, in turn, directs subsequent patterning stages of Ato (Li and Baker, 2001). The enhancement of IGs during stage-2 features two discrete modes: early stage-2 presumed to be driven by *ato-3’*, and late stage-2 driven primarily by *5’-ato* (Jarman et al., 1994; Sun et al., 1998; Majot and Bidwai, 2017). In addition to its neurogenic role, N signaling also represses Ato, facilitating R8 individuation. This repressive role for N utilizes the transcription factor Su(H), which elicits the expression of bHLH repressors of the *E(spl)* locus (Bailey and Posakony, 1995; Jennings et al., 1994). Previous studies demonstrate that the proneural N signal occurs independently of Su(H) (Li and Baker, 2001). E(spl) is required for the onset of stage-3 Ato patterning, in which E(spl) antagonize Ato via the 5’ enhancer (Majot and Bidwai, 2017).

In addition to N, EGFR plays an active role in Ato patterning. The EGFR signal is transduced to the nucleus via activated (di-phosphorylated) MAPK. Indeed, MAPK is strongly activated within IGs, with peak activity occurring during late stage-2 through stage-3, spanning the time of onset of Ato repression (Chen and Chien, 1999; Kumar et al., 2003). Additionally, MAPK activation is cell-autonomously dependent upon Ato, and may, therefore, be dependent upon N signaling as well (Chen and Chien, 1999; Lim and Choi, 2004). Thus, the correlation between MAPK activity and Ato repression bears further investigation for possible nodes of N-EGFR crosstalk with regard to Ato expression.

To date, only E(spl)M8 (referred to as M8) has emerged as an apparent genetic link between N and EGFR-MAPK (Bandyopadhyay et al., 2016, Majot et al., 2015). Both genetic and biochemical evidence suggest that M8 directly antagonizes Ato via its 5’ enhancer following the formation of IGs (Majot and Bidwai, 2017, Karandikar et al., 2004, Kahali, et al, 2010). N and Su(H) are required for the expression of M8 and genetic evidence suggests that MAPK phosphorylates the C-terminal domain of M8 to potentiate M8’s repression of Ato (Bandyopadhyay et al., 2016; Bose et al., 2014). However, M8 fails to account for the entirety of cooperativity between N and EGFR, as evidenced by investigations with the dominant M8 allele, *E(spl)^D^*. *E(spl)^D^* encodes a truncated protein variant which lacks the regulatory C-terminal domain, producing a constitutively active form of M8 (Tietze et al., 1992; Nagel et al., 1999; Kahali et al., 2010). Despite being hypermorphic, *E(spl)^D^* alone elicits only minor morphological eye defects as compared to mutations that perturb N or EGFR signaling (Majot and Bidwai, 2017; Li et al., 2003; Lesokhin et al., 1999). Thus, if MAPK is required for the repression of Ato, additional targets than M8 likely exist.

Anterior open (Aop, also known as Pokkuri or Yan), an ortholog of human TEL1/ETV6, may also link N and EGFR signaling during R8 specification. Aop, an ETS repressor, is expressed throughout a variety of developmental contexts (Rebay and Rubin, 1995). In its native state, Aop represses target genes through direct enhancer binding. However, repressor capacity is attenuated upon phosphorylation by MAPK (Rebay and Rubin, 1995; O’Neill et al., 1994). In eye development, Aop is first expressed within the MF and is present in many cells of uncommitted fate (Boisclaire-LaChance et al., 2014; Olson et al., 2011). Previous investigations have revealed several roles for Aop in retinogenic cell fate decisions (Tei et al., 1992; Lai and Rubin, 1992; Rogge et al., 1995). *aop* flies were originally characterized with respect to eye development for having supernumerary R7 photoreceptors, which are produced in a process that was later shown to be both N and Ras-MAPK-dependent (Tei et al., 1992; Tomlinson et al., 2011). Although the loss of Aop disrupts R8 specification, the reason for such disruption has remained unclear (Rogge et al., 1995; Olson et al., 2011). Aop has been shown to regulate Armadillo (Arm, orthologous to β-CATENIN), and potentially, by association, Wg/Wnt signaling (Olson et al., 2011; Caviglia and Luschnig, 2013). However, the links between Aop and R8 patterning have remained elusive.

In this study, we have probed Aop in its relationship to Ato patterning. We reveal that Aop may constitute a significant node for cross-regulation of eye development through both N and EGFR-MAPK signaling given that Aop requires Su(H) for its eye-specific expression, and that its activity is lost upon phosphorylation by MAPK (Rohrbaugh et al., 2002; O’Neill et al., 1994; Rebay and Rubin, 1995). We demonstrate that Aop is coexpressed with Ato only during early stage-2 of Ato patterning, during the initial moments of IG formation. The removal of aop prevents proper Ato patterning, and greatly diminishes R8 formation. We further provide evidence that both the Su(H)-independent and Su(H)-dependent mechanisms of N signaling occur concurrently with one another in the MF, rather than sequentially. Su(H) is required to induce Aop during Ato’s inductive stages, and genetic interactions suggest that Aop functions at this time to block E(spl) activity against Ato. This study provides several lines of evidence to facilitate deeper investigation into crosstalk between N, EGFR, and their regulation of Ato’s discrete enhancers.

## Results

### Aop cell-autonomously regulates Ato

To explore whether Aop regulates *ato*, we observed Ato expression in mitotic clones that were derived using the FLP-FRT recombination system (Xu and Rubin, 1993). *aop* clones feature a striking, novel Ato patterning defect in which Ato is present, but with reduced range when compared to WT (Fig. 2A,B). Despite seemingly normal initiation at the anterior margin of its expression domain, Ato fails to upregulate into IGs. Despite the Ato deficiency, some Ato-positive individuals emerge from the MF and likely develop into R8s (Fig. 2A, yellow arrow), as indicated by Sens immunolabeling that shows occasional *aop* mutant R8s (Fig. 2C,D; Olson et al., 2011). Furthermore, these Ato-positive individuals are not likely to be re-emergent Ato expression, but a continuation of Ato expression from the MF, as Fig. 2B (white arrowheads) shows that high-level Ato-positive individuals are observed in the anteroposterior position where IGs normally form. Importantly, the boundary between WT and mutant cells indicates that Aop cell-autonomously regulates IG maturation, as Ato remains unaffected in WT border cells while such expression is reduced in neighboring mutants (Fig. 2B, yellow arrow). Mitotic clones of *aop^yan1^*, a mutant that elicits less severe adult phenotypes than those of *aop^1^*, were similarly assayed (Fig. 2E). In *aop^yan1^*, Ato was weaker and accompanied by patterning deficiencies that were similar, though much less severe, to those of *aop^1^*.

**Figure 2.**
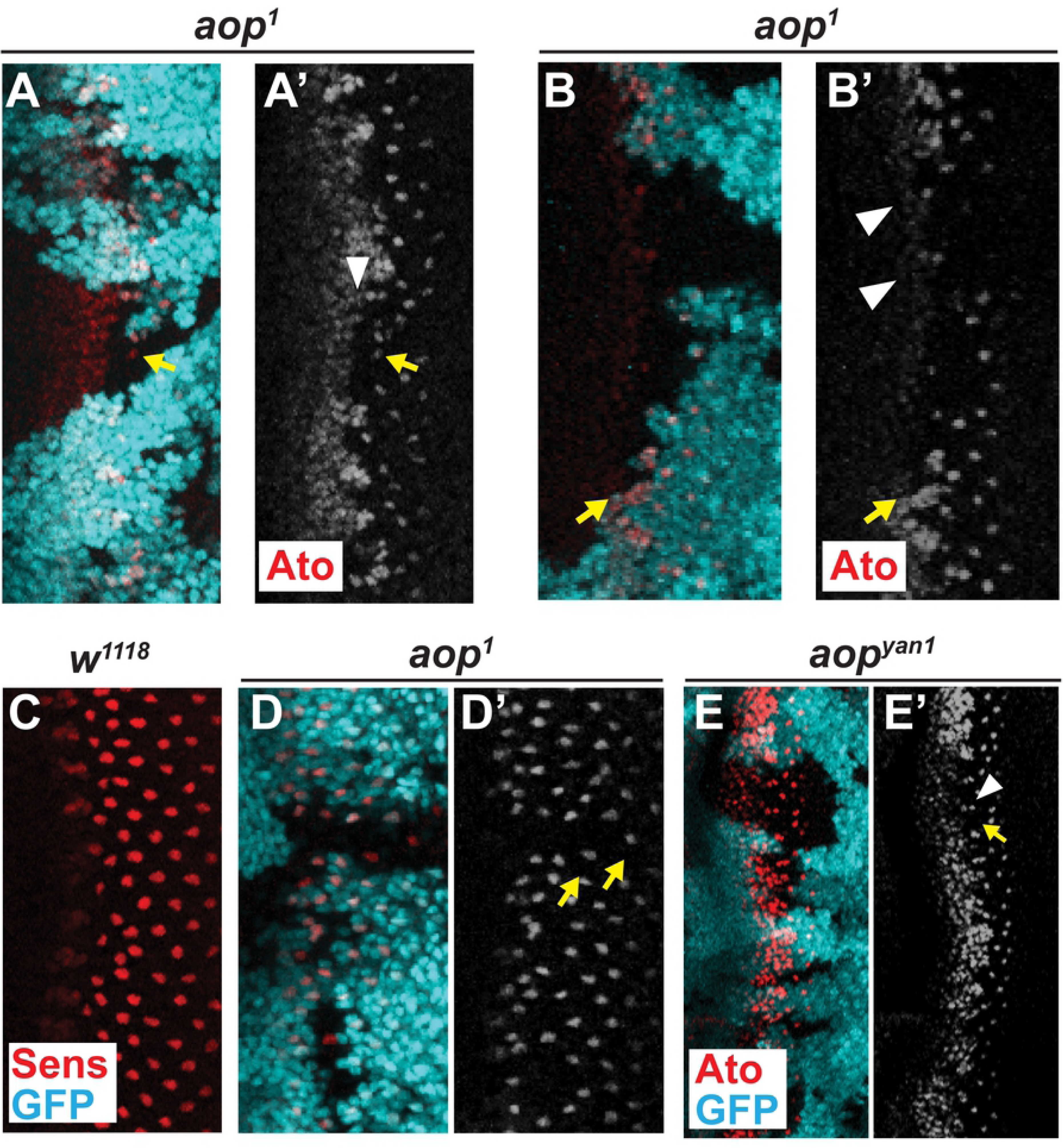
Aop cell-autonomously regulates Ato. (A) *aop* mutants are Ato-deficient. GFP (cyan) indicates heterozygous or WT tissue. Despite initiating as in WT, Ato (red) fails to form mature IGs (arrowhead), and isolated *aop* mutants occasionally maintain Ato (arrow). (B) Isolated Ato^+^ mutants likely maintain their expression from the MF (arrowheads), demonstrating that *aop* is required cell-autonomously for Ato IG formation (arrow). © Sens expression initiates in IGs and resolves to R8s in WT, as shown. (D) *aop^1^* mutants lack Sens expression, with the exception of occasional individuals (arrow). (E) *aop^yan1^* mutants feature diminished Ato within IGs (arrowhead) and some missing R8s (arrow). *aop^yan1^* is a weaker loss-of-function allele than *aop^1^*. Genotypes: (A,B,D) *aop^1^ frt40A*/*ubiGFPnls frt40A*;*eyFLP*/+,(C) *w^1118^*, (E) *aop^yan1^ frt40A*/*ubiGFPnls frt40A*;*eyFLP*/+.

### *aop* clones do not affect the timing of Ci processing or Dpp expression

Next, we assessed whether the Ato patterning defects in *aop* mutants are attributable to misregulation of the morphogenic pathways Hedgehog (Hh) and Decapentaplegic (Dpp; Fig. 3). In WT, Hh is secreted from differentiating photoreceptors and drives anterior progression of the MF (Ma et al., 1993). Cells of the MF respond to Hh through the accumulation of the active transcription factor, Cubitus interruptus (Ci^ACT^; Fig. 3A; Motzny and Holmgren, 1995). Once the MF passes, Ci^ACT^ is abruptly lost from differentiating photoreceptors (Fig. 3A). In *aop* cells, Ci^ACT^ accumulates ahead of the MF as is WT and is abruptly lost with the passage of the MF, suggesting that Hh signal response is largely unperturbed (Fig. 3B, arrows).

**Figure 3.**
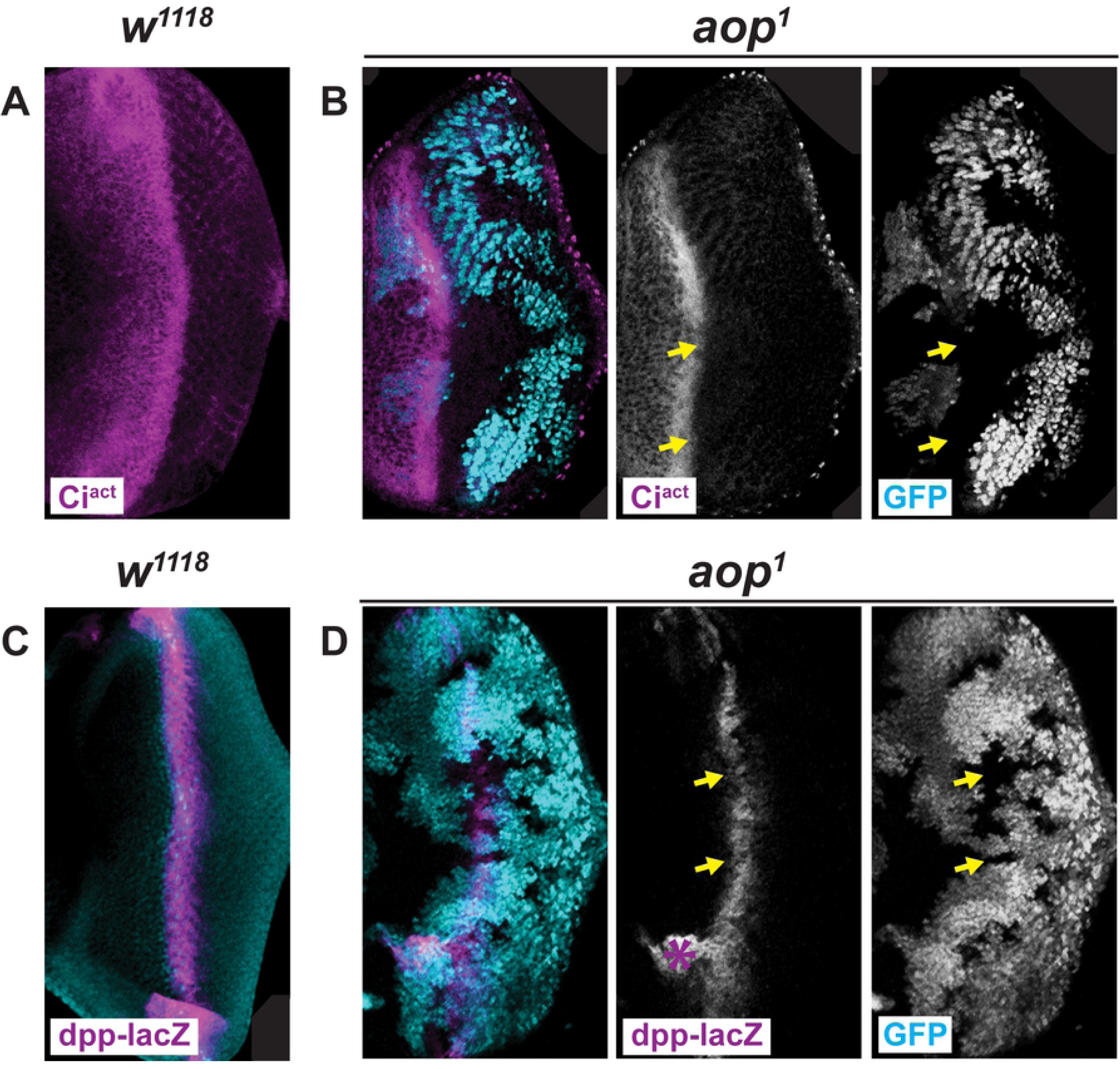
*aop* clones do not affect the timing of Ci processing or Dpp expression. (A) Hh-signaling response is dictated by the loss of a repressor form of Ci (Ci^rep^) in favor of the accumulation of the activator form (Ci^act^). Ci^act^ accumulates most strongly anterior to and within the MF and is absent from differentiating retinulae. (B) *aop* mutants do not disrupt the timing nor apparent strength of the Hh response as monitored by Ci^act^. (C) Dpp, which is resultant of Hh signaling, is expressed in cells of the MF. (D) *dpp-lacZ*, often used as an analog for Dpp expression, is expressed in *aop* mutants, suggesting that the Hh response occurs as in WT. Genotypes: (A) *w^1118^*, (B) *aop^1^ frt40A*/*ubiGFPnls frt40A*;*eyFLP*/+, (C) P{*dpp-lacZ*}/+ (chr. III), (D) *aop^1^ frt40A*/*ubiGFPnls frt40A*;*eyFLP*/P{*dpp-lacZ*}.

The morphogen Dpp is also employed in eye development. Hh stimulates the secretion of Dpp from WT cells of the MF (Heberlein et al., 1993). Thus, Dpp expression can be used as an assessment of the normality of the gene expression program within the MF. Using the *dpp-lacZ* reporter, which reports from a dorsoventral band of cells that approximately comprise the MF (Fig. 3C; Blackman et al., 1991), we confirm that loss of *aop* has negligible effect on *dpp* report (Fig. 3D, arrows). When taken together, the Ci^ACT^ and *dpp-lacZ* results demonstrate that the low level Ato observed in *aop* clones does not reflect a loss of the morphogenic signaling from the Hh and Dpp pathways within the MF. In combination with the Ato phenotype of Fig. 2, these results indicate that Aop likely participates in IG formation and not initial Ato induction.

### Ato and Aop colocalize during IG formation

*aop* mutants generally fail to differentiate into R8s (Rogge et al., 1995; Olson et al., 2011), as evidenced by Senseless (Sens), an R8 differentiation marker that, in the retina, is preceded and directed by the expression of Ato (Fig. 2D; Frankfort et al., 2001). In WT, Sens initiates in small clusters within IGs and, along with Ato, are resolved to individuated R8s. We have now shown that IG formation exhibits cell-autonomous dependence on Aop (Fig. 2). Thus, we assessed the coexpression of Aop and Ato. Aop is first observed in the MF at the same approximate time as Ato (Fig. 4). Ato and Aop both progress through staged expression patterns. During Ato’s stage-1, Aop is absent. When Ato is first organized into IGs, Aop colocalizes with Ato (Fig. 4B, white arrows). Later, as IGs become more pronounced, Aop is lost (Fig. 4A, arrows; Fig. 4B, yellow arrows), and does not colocalize with Ato throughout the posterior of the eye.

**Figure 4.**
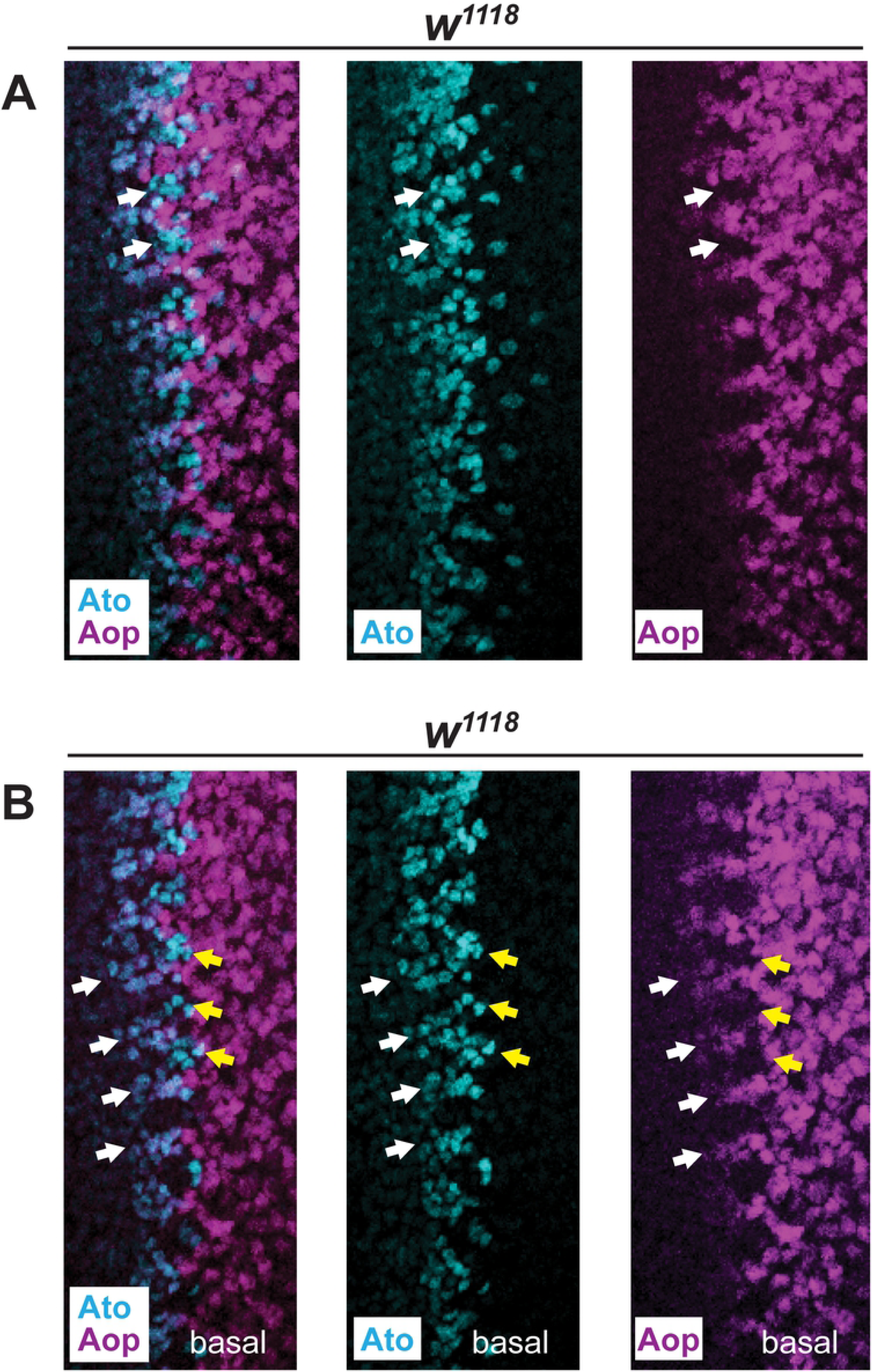
Ato and Aop colocalize during IG formation. (A) Ato and Aop are both expressed in the developing retina starting in the MF. Mature IGs, marked by arrows, lack Aop. (B) A basal section of the same tissue shown in A demonstrates the colocalization of Ato and Aop where IGs are forming (white arrows). Yellow arrows indicate Ato^+^ mature IGs, similar to those highlighted in B. Genotypes: (A,B) *w^1118^*.

### Aop is induced by N signaling during Ato patterning

We next assessed Aop’s role within the context of MF-related N signaling. Ato elicits N signaling, and subsequently drives Su(H)-responsive genes, thus we probed Aop expression in *ato*, *N* and *Su(H)* backgrounds (Fig. 5; Baonza and Freeman, 2001). Aop is expressed in *ato* mutants (Fig. 5A) but its expression is delayed, similar to expression of E(spl) in proneural mutants (Lim et al., 2008). Additionally, in *ato* mutants, Aop immunostain is observed with less intensity as in WT (Fig. 5A). In *N* mutants, however, Aop fails to express in either the MF or the posterior eye disc (Fig. 5B). Similarly, *Su(H)* mutants lack Aop labeling throughout the eye disc (Fig. 5C). In addition to the cell-autonomous absence from the MF in *Su(H)* mutants (Fig. 5C, arrowhead), Aop is lost from WT cells that neighbor the mutant tissue toward the posterior of the eye disc (Fig. 5C, arrows) suggesting a shift in Su(H)-dependent regulation as retinal patterning progresses past the MF.

**Figure 5.**
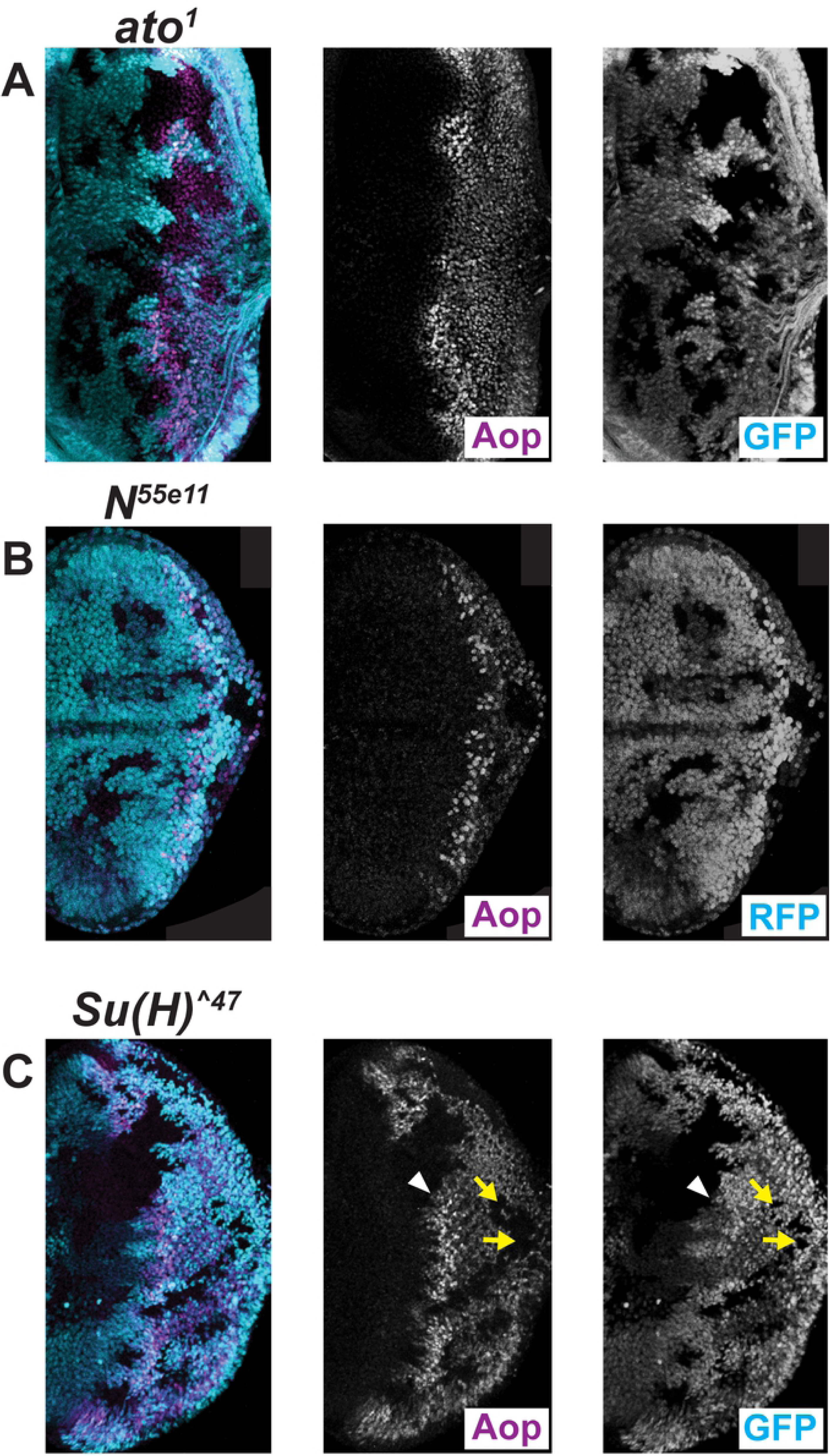
Aop is induced by N signaling during Ato patterning. (A) *ato* mutant cells exhibit non-autonomous, delayed Aop expression. Overall, *ato^1^* cells fail to properly induce Aop as early occurs in WT. (B) Removal of Notch elicits cell-autonomous loss of α-Aop immunostaining. (C) In the absence of Su(H), α-Aop labeling is not detected. Near the MF, Aop cell-autonomously requires Su(H) (arrowhead), whereas toward the posterior of the eye disc, Aop is also absent from WT cells that border mutant regions (arrows). Genotypes: (A) *frt82B ato^1^*/*frt82B ubiGFP eyFLP*, (B) *N^55e11^ frt19A*/*ubiRFP hsFLP frt19A*, (C) *Su(H)^^47^ frt40A P{l(2)Bg35+}*/*ubiGFPnls frt40A*;*eyFLP*/+.

### Armadillo is upregulated in neurogenic backgrounds

Past reports indicate that Aop regulates Arm, which accumulates more quickly in the adherens junctions of *aop* mutants than in WT tissue (Fig. 6; Olson et al., 2011). Olson et al. proposed that the Arm enhancement was disruptive to neurogenesis through putative enhancement of Wg signaling. However, we find that Arm accumulation coincides with neural hypertrophy in multiple mutant backgrounds. As demonstrated in other *aop* mutants, *aop^1^* features Arm enhancement (Fig. 6B). To note, *Su(H)* mutants, which also lack Aop (Fig. 5C; Rohrbaugh et al., 2002), feature both Arm accumulation (Fig. 6C) and neural hypertrophy (Li and Baker, 2001). Similarly, *E(spl)* clones also feature both Arm accumulation (Fig. 6D) and neural hypertrophy. Thus, Arm accumulation does not preclude the possibility of neural hypertrophy, as previously suggested.

**Figure 6.**
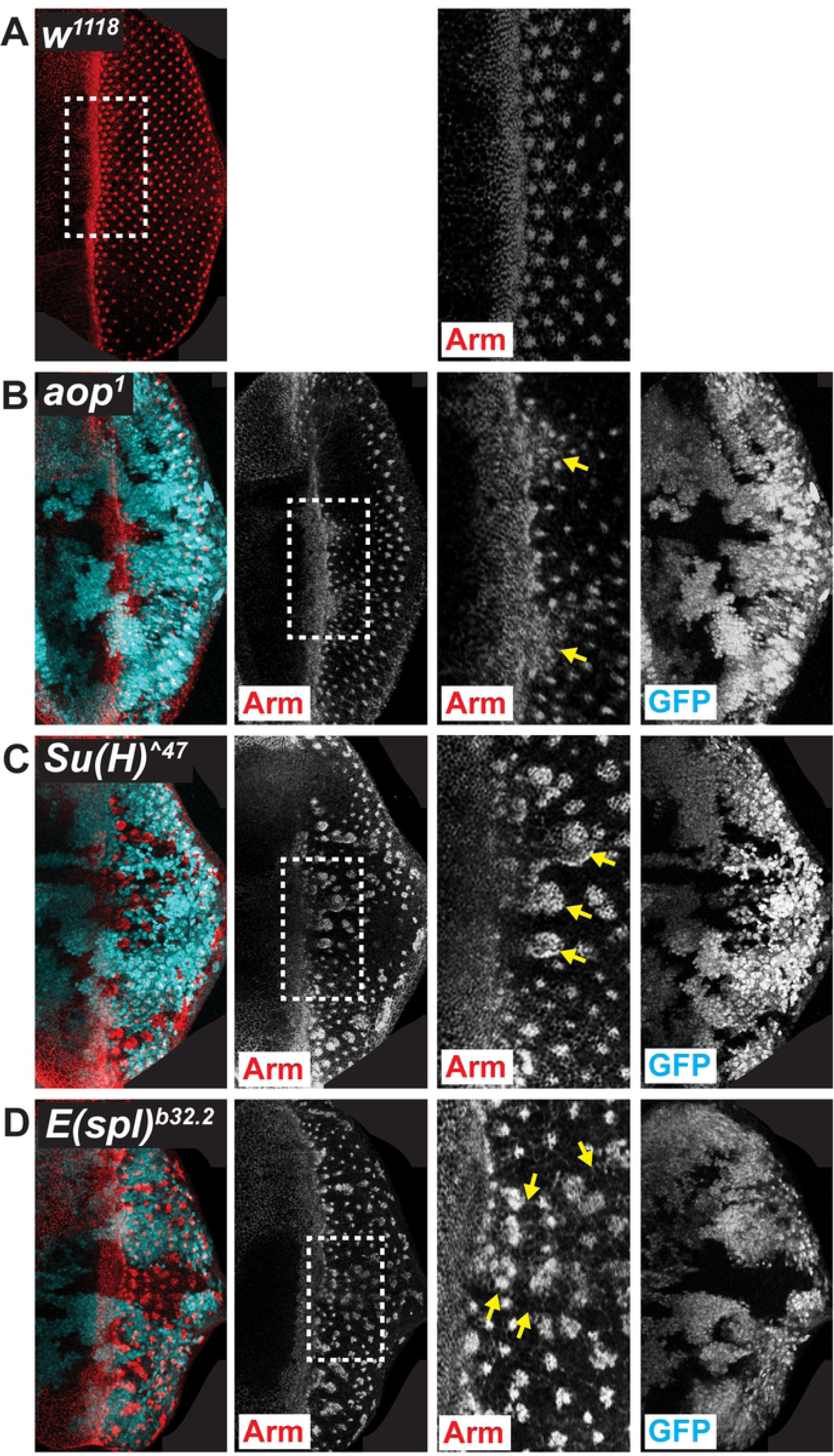
Armadillo is upregulated in neurogenic backgrounds. (A) Arm (red) accumulates at adherens junctions, which are most highly concentrated within the MF and rosettes. Anterior is left. (B) *aop* mutant clones accumulate Arm. As for all clonal analyses, far left panel is merge, followed to the right by individual channels. Mutant tissues are marked by the absence of GFP. Inset at far right shows continued apical constriction and corresponding Arm accumulation of *aop* mutants (arrows). (C,D) *Su(H)* and *E(spl)* mutants, respectively, display similar Arm enhancement phenotypes. Arrows in (C) indicate several clusters of persistent apical constriction. Arrows in D delineate a large *E(spl)* clone. Note the persistent high-level background staining in *E(spl)* clones. Genotypes: (A) *w^1118^*, (B) *aop^1^ frt40A*/*ubiGFPnls frt40A*;*eyFLP*/+, (C) *Su(H)^^47^ frt40A P{l(2)Bg35+}*/*ubiGFPnls frt40A*;*eyFLP*/+; (D) *frt82B Df(3R)E(spl)^b32.2^ P{gro^+^}*/*frt82B ubiGFP eyFLP*.

### Aop is required for proper expression of both *ato* enhancers

We further dissected Aop regulation of Ato through use of *ato-3’* and *5’-ato* enhancer reporters (Fig. 7; Sun et al., 1998). Enhancer activity was assessed in the *aop^1^*/*aop^yan1^* background in which adults display aberrant ommatidial patterning (Rogge et al., 1995). Despite adult pattern defects, Ato is observed within the MF of transheterozygous animals, but at low levels and lacking consistent IG formation. Regardless of the observed Ato defect, R8s consistently form and are labeled by Sens (Fig. 7B,D). In WT, *5’-ato* initiates report in IGs and can be observed throughout the posterior of the eye disc (Fig. 7A). In *aop* flies, *5’-ato* report initiates where IGs should normally form, but only at low levels that lack discernable clusters (Fig. 7B). WT *ato-3’* reports strongly within the MF and resultant immunostain tapers toward the posterior eye disc (Fig. 7C). In *aop* flies, *ato-3’* can be detected in the MF, but at greatly muted levels (Fig. 7D). The antennal expression of either enhancer reporter was unaffected in *aop^1^*/*aop^yan1^* animals (not shown).

**Figure 7.**
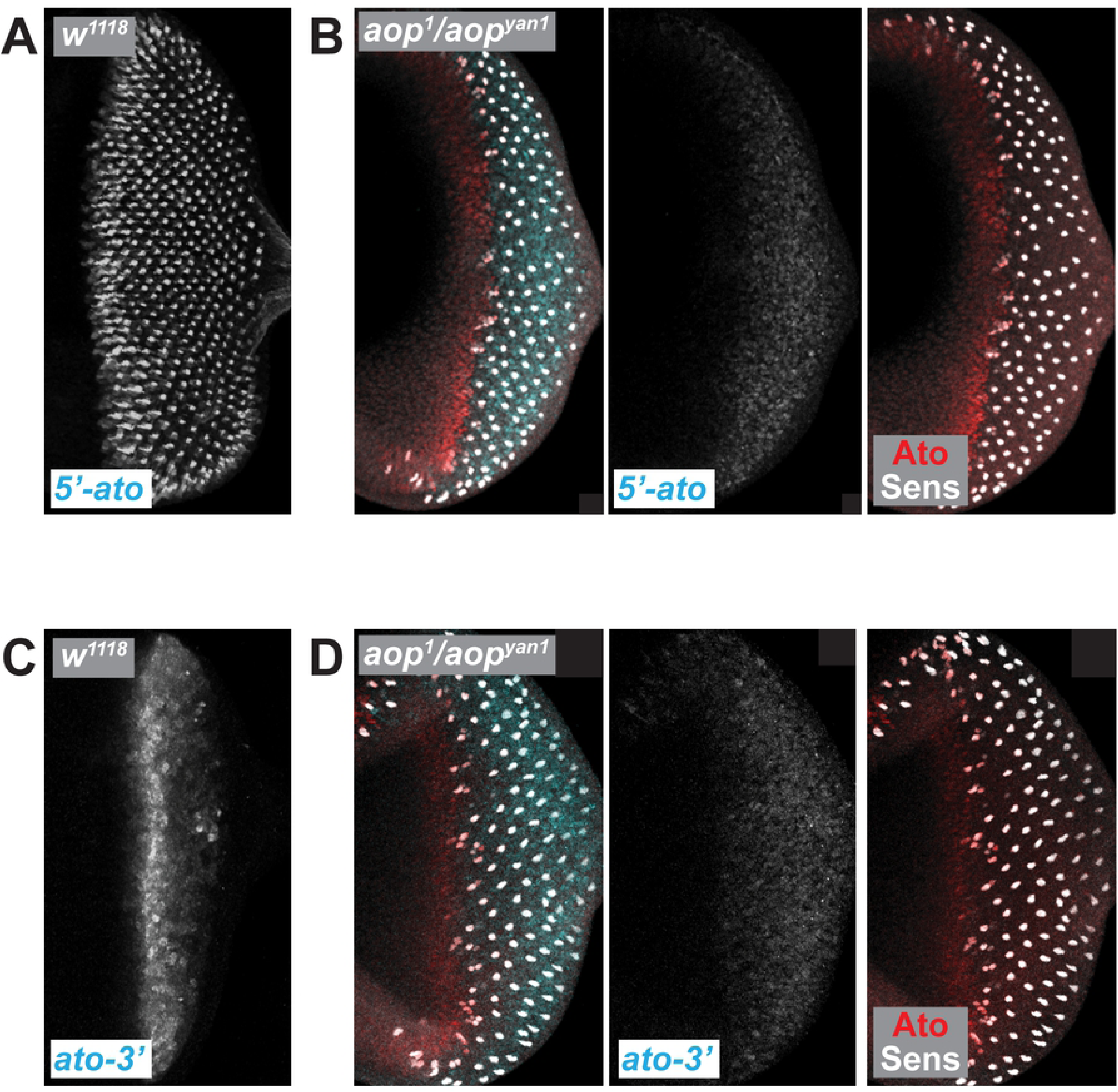
Aop is required for proper expression of both *ato* enhancers. (A) In WT, *5’-ato-lacZ* reports in IGs and the signal perdures distinctly in R8s. (B) *aop* mutants feature greatly perturbed *5’-ato* activity. (C) In WT, *ato-3’-lacZ* reports in cells of the MF. (D) *aop* mutants elicit severe, but incomplete loss of *ato-3’* activity. In (B,D), Ato (red) and Sens (gray) are shown to demonstrate position of the MF. Genotypes: (A) P{*w^+mC^ ato5’F:9.3*}/+, (B) *aop^1^ frt40A*/aop^yan1^ P{*w^+mC^ ato5’F:9.3*}, (C) P{*w^+mC^ ato3’F:5.8*}/+, (D) *aop^1^ frt40A*/*aop^yan1^* P{*w^+mC^ ato3’F:5.8*}.

### Aop does not repress E(spl)

Having established that Aop is required for IG formation, we next assessed whether Aop might regulate the timing of E(spl) expression. In WT, E(spl) are expressed dynamically across the MF (Fig. 8A; Baker et al., 1996; Majot and Bidwai, 2017). In *aop* clones, E(spl) immunolabeling resembles WT (Fig. 8B). Although fine-scale aberrations in E(spl) expression may occur, no clear effect to overall patterning or timing of expression (precocious or delayed) are observed in *aop* mutants (Fig. 8B). To more directly test the possibility that Aop represses E(spl), we assessed the effects of force-expressed *aop^act^*. *aop^act^* encodes a constitutively active form of Aop wherein the MAPK-target Ser residues are mutated to Ala to prevent inhibition of Aop by MAPK (Rebay and Rubin, 1995). In wandering third instar larvae, the *dppGAL4* driver produces GAL4 in developing antennae along the dorsal and ventral margins and in a swath of cells that stretch between the two margins across the center of the antennal disc (Fig. 8C; Blackman et al., 1991). As shown using a UAS-GFP reporter, GAL4 is produced in a cluster of cells along the dorsal margin that overlaps with cells that express E(spl) (Fig. 8C). If Aop binds and represses genes of the *E(spl)* locus through *trans* interaction, reduced E(spl) expression would be observed in the presence of exogeneous Aop^act^. However, overexpression of Aop^act^ resulted in an enlarged E(spl) expression domain, as was similarly observed with that for Ato and Sens (Fig. 8D).

**Figure 8.**
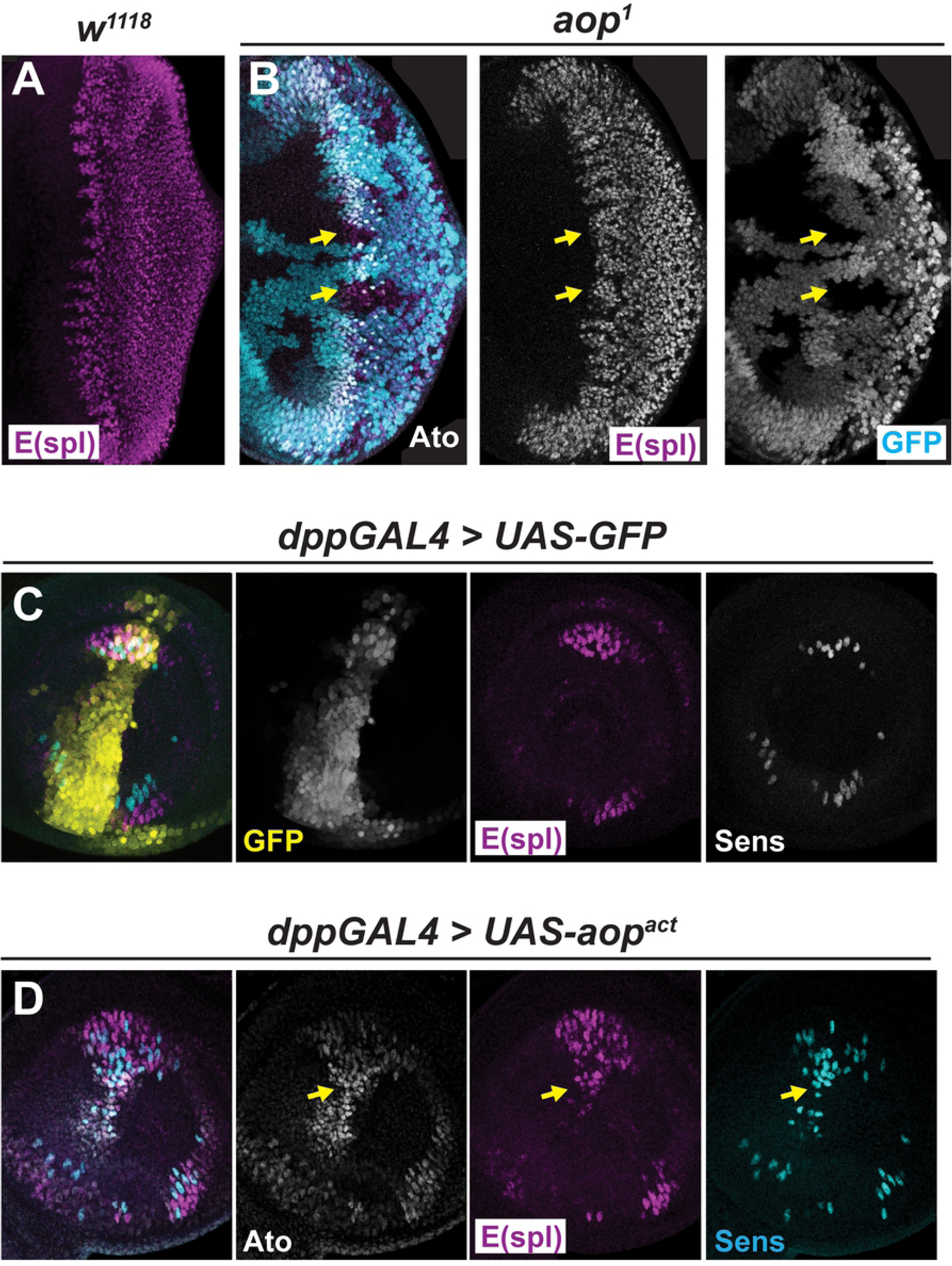
Aop does not repress E(spl). In WT, Ato and E(spl) co-localize in early-stage Ato+ cells (Majot and Bidwai, 2017), but not in mature IGs (yellow arrowhead). (A) A representative pattern of WT E(spl) expression. (B) *aop* mutants fail to perturb E(spl) (arrows). (C) dppGAL4 expression in antennal discs of a third instar larva. GFP co-localizes with E(spl) and Sens in several regions at the periphery of the tissue. (D) Aop^act^ fails to repress E(spl) from within cells of the dppGAL4 expression domain. Sens and Ato are also present in expanded expression patterns. Genotypes: (A) *w^1118^*, (B) *aop^1^ frt40A*/*ubiGFPnls frt40A*;*eyFLP*/+, (C) *P{UAS-GFP-nls}*/+;*P{GAL4-dpp-blk1*}/+, (D) *P{w^+mc^ UAS-aop^act^}*/+;*P{GAL4-dpp-blk1}*/+.

### Aop sensitizes Ato to repression by E(spl)

Although Aop has little effect on E(spl) expression within the MF, colocalization of Ato, E(spl), and Aop^act^ in the antennal disc suggests that WT R8 specification may require Aop through an indirect effect on E(spl) activity toward Ato. Thus, we observed Ato and Sens expression in clones dually mutant for *aop* and *E(spl)* (Fig. 9A). Removal of *E(spl)* alone elicits widespread neurogenesis and upregulation of both Ato and Sens (Ligoxygakis et al., 1998). Together, *aop* and *E(spl)* elicit a similar, though weaker, phenotype in which Ato and Sens are expressed in many cells of the MF. As occurs in *E(spl)*, Ato wanes toward the posterior of the eye although Sens remains strong throughout (Fig. 9A; Li and Baker, 2001).

**Figure 9.**
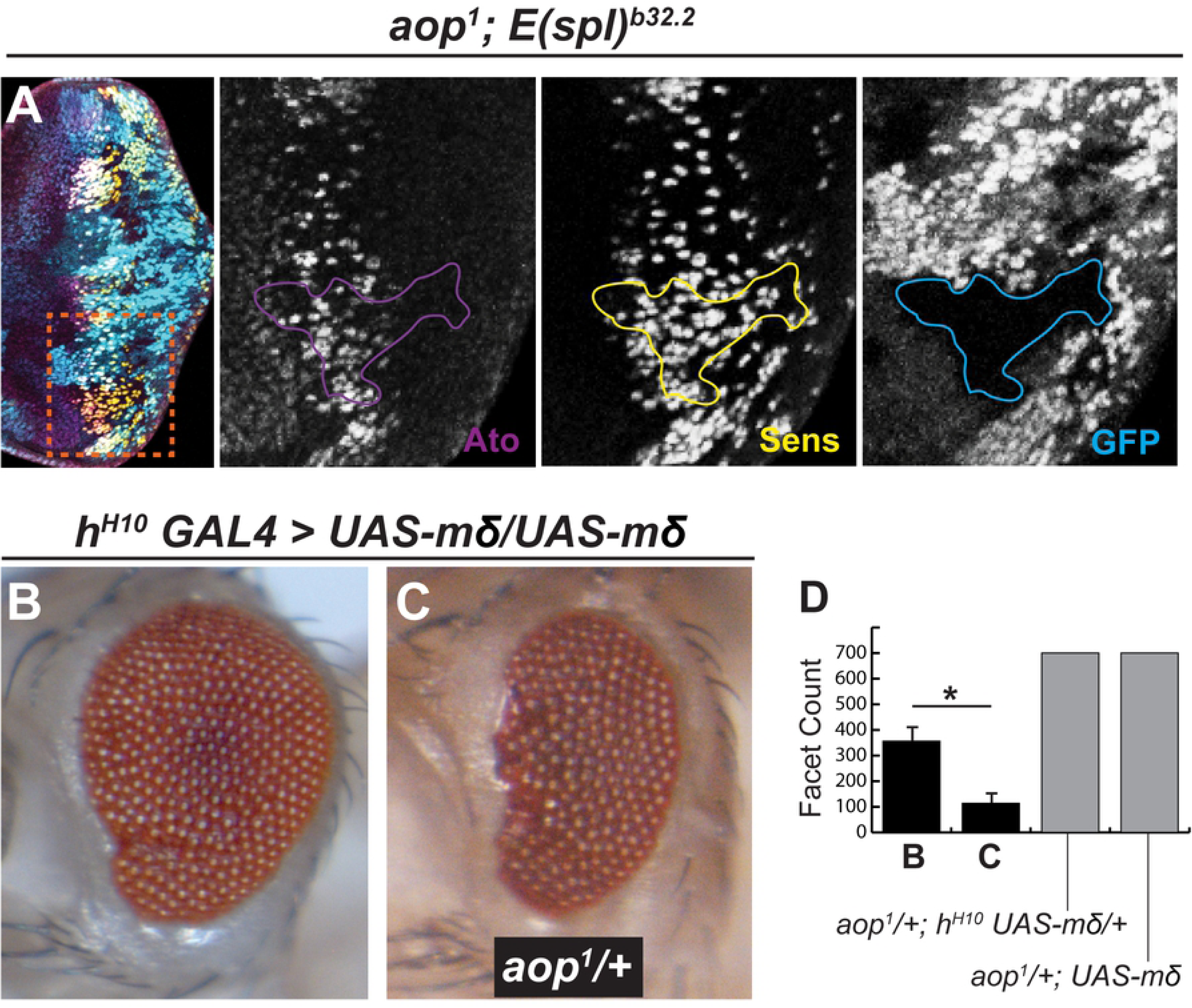
Aop sensitizes Ato to repression by E(spl). (A) *aop*, *E(spl)* dual mutant clones feature α-Ato (magenta) and α-Sens (gold) staining, as opposed to reduced α-Ato in *aop* clones (Fig. 2). Insets focus on large dual mutant clone, note sporadic though prevalent Ato and broad expression of Sens, indicating that most aop, E(spl) mutants are specified as R8s. (B) *h^H10^ GAL4* driving two copies of transgenic *UAS-E(spl)mδ*. The eye is reduced, mispatterned and features a slight cleft at the anterior margin. (C) The introduction of *aop* loss-of-function to the genotype in B further reduces the eye. Quantitation to the right of C; n≥10, asterisk (*) indicates P<0.001. Genotypes: (A) *aop^1^ frt40A*/*ubiGFPnls frt40A;frt82B Df(3R)E(spl)^b32.2^ P{ry^+t7.2^=gro}/frt82B ubiGFP eyFLP* (B) *h^H10^ GAL4 P{w^+mc^ UAS-E(spl)mδ}*/*P{w^+mc^ UAS-E(spl)mδ}*, (C) *aop^1^ frt40A*/+; *h^H10^ GAL4 P{w^+mc^ UAS-E(spl)mδ}*/*P{w^+mc^ UAS-E(spl)mδ}*.

Having demonstrated that *aop* sensitizes *ato* to E(spl), we assessed whether *aop* enhances eye defects elicited by E(spl) gain-of-function. The *hairy^H10^* enhancer trap (*h^H10^ GAL4*) drives GAL4 expression anterior to and within the MF (Ellis et al., 1994). No overt eye defects (see below) are intrinsic to *aop*^*1*/+^; *UAS-mδ* flies on their own, or upon *h^H10^ GAL4* driven expression of a single copy of *E(spl)mδ* in an *aop*^*1*/+^ background (Fig. 9D). However, forced-expression of two copies of *E(spl)mδ* driven by *h^H10^ GAL4* elicits aberrant Ato immunolabeling, a reduced eye, and frequently, an anterior cleft (Ligoxygakis et al., 1998) that resembles the furrow-stop phenotype (Chanut et al., 2000). *aop* significantly enhances the eye defect of force-expressed *E(spl)mδ*, consistent with a role in further enhancing *ato* repression (Fig. 9B,C,D).

## Discussion

In the developing retina, Aop is a downstream component of the N signaling pathway. Previous efforts have identified the presence of Su(H) paired-sites (a strong predictor of Su(H) binding) in an upstream enhancer of Aop (Rohrbaugh et al., 2002). Enhancer-reporter constructs were able to clarify the importance of such sites in regard to eye-disc specific expression of Aop. Additionally, immunochemical approaches have verified that in the absence of Su(H), Aop is not expressed in either the MF (Fig. 5C) or throughout the eye disc (Rohrbaugh et al., 2002). In the MF, Ato induces N signaling (Baonza and Freeman, 2001). As such, we also demonstrate that timely induction of Aop requires the Ato-N-Su(H) signaling axis (Fig. 5). Ato exhibits non-autonomous control of Aop, likely the result of N signaling that is properly initiated from nearby adjacent WT cells. A similar observation was noted in *da* clones when immunostained for E(spl) (Lim et al. 2008), suggesting that this expression phenotype may be typical of Su(H)-respondent genes in the MF.

This study is the first to have explored Aop in the context of proneural gene regulation. Analysis of *aop* clones reveals a distinct Ato phenotype that has not yet been documented: initial, stage-1, Ato remains unaffected while later Ato stages are nearly absent. Despite the presence of occasional Ato+ individual cells as shown in Fig. 2, IGs are never observed in *aop* tissue in >30 animals that have been analyzed. Furthermore, this phenotype is definitively cell-autonomous, as evidenced by clonal boundaries that cross through IGs (Fig. 2B). In such cases, WT cells feature the normal appearance of IGs whereas mutant cells are sharply contrasted by Ato deficiency. In order to exert its cell-autonomous maintenance of Ato expression, Aop must be present during or before the point at which Ato is lost from *aop* mutants. Accordingly, Aop colocalizes with Ato only during early stage-2, where IG formation initiates (Majot and Bidwai, 2017). By late stage-2, Aop, as with other Su(H)-responsive genes, is absent from IGs. Thus, early stage-2 is the only appropriate window for Aop to exert its neuroprotective function toward Ato and R8 fate.

In light of the stage-specific *aop* phenotype, we reasoned that there might be an enhancer-specific effect on *ato*. Unsurprisingly, *5’-ato* report was highly aberrant in *aop* mutant eyes. *ato-3’* was also affected such that although the enhancer report initiated appropriately, it failed to build to the same high levels detected in WT eyes. Given that Aop colocalizes with Ato during early stage-2 (when Ato is solely derived from *ato-3’*) and that *ato-3’* fails to strongly report from *aop* mutants, it is plausible that Aop exerts its neuroprotective effect toward *ato-3’* concurrently with E(spl) repression of *5’-ato*. Thus, during early IG formation while E(spl) actively repress *5’-ato* (Majot and Bidwai, 2017), Aop may function in parallel to indirectly promote continued *ato-3’* induction. As shown with clonal analysis and force-expression assays, Aop does not exert any notable influence on the expression of E(spl). However, E(spl) is functionally sensitive to *aop* in the MF, as demonstrated by *aop*-mediated enhancement of a classic E(spl) gain-of-function phenotype (Fig 9).

Stage-2 is a critical inflection point in Ato patterning. At this time, Ato successfully switches from strict *ato-3’* dependence to primary dependence upon *5’-ato* (Fig 10). As discussed previously, stage-2 is best considered as divided into discrete sub-stages. Early stage-2 is defined by the role of E(spl) (Majot and Bidwai, 2017). As observed with Aop, E(spl) are expressed at the onset of IG formation and the absence of E(spl) allow precocious activation of *5’-ato*, indicating that during this time, *ato-3’* is responsible for the WT production of Ato while *5’-ato* is primed for expression. Late stage-2 is defined by the absence of E(spl) and Aop from these cells, partly through function of the Su(H)-repressor Roughened eye/Roe (Melicharek et al., 2008; del Alamo and Mlodzik, 2008; Majot and Bidwai, 2017). Stages-3 and -4 are characterized by the return of E(spl). However, Ato is at this point repressed from all cells except those that are to become R8s. Notably, at stages-3 and -4, the EGFR-MAPK pathway is active. EGFR-MAPK has been demonstrated to repress the function (and possibly expression) of Aop. It is not clear from our data or past studies whether Aop’s absence from stage-3 and -4 Ato IGs is the result of EGFR-MAPK activity or some other mechanism.

**Figure 10.**
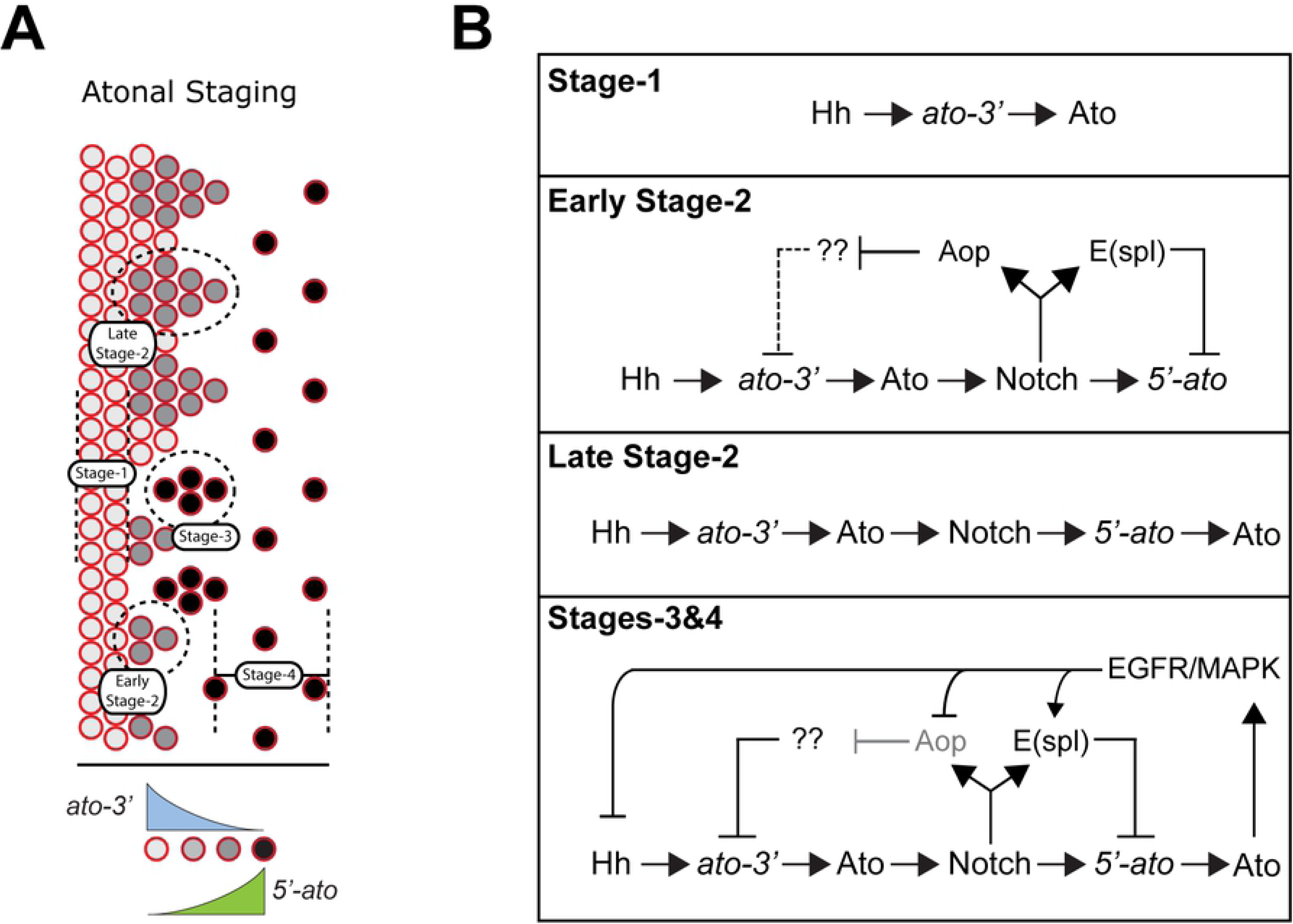
Aop is required during early stage-2. (A) Ato progresses through distinct stages of its expression pattern, stages-1 through -4, from a broad profile to individuated R8s. (B) Stage-1 Ato is driven by non-autonomous, morphogenic Hh signaling. Early stage-2 is characterized by the activation of the Notch pathway and the expression of Su(H)-respondent genes Aop and E(spl). In the absence of Aop, E(spl) is capable of repressing Ato in early stage-2. Late stage-2 features the transient repression of some, if not all, Su(H)-respondent genes. This step is notably observed as increased intensity of Ato immunostaining within IGs. Stages-3 and -4 see both the return of E(spl) and loss of Ato in all cells but the presumptive R8s. It remains unclear whether EGFR/MAPK activity at this stage affects Aop expression, as Aop immunostain is poor to nonexistent in stage-3, -4 IGs.

Thus, data support a mechanism of Ato regulation in which Aop and E(spl) respectively regulate *ato-3’* and *5’-ato* in parallel (Fig. 10B). In the absence of MAPK activity, Aop maintains a protective function toward the 3’ enhancer. However, in the presence of active MAPK, Aop would be relieved of this duty. In support of this mechanism, *in situ* hybridization against *ato-3’* reveals that the 3’ enhancer is restricted into clusters that resemble early stage-2 IGs prior to loss of enhancer activity (Sun et al. 1988). Consistent with this, loss of both E(spl) and Aop leads to chaotic upregulation of Ato and pan-R8 neurogenesis, similar to that observed in E(spl)-only mutants.

Thus, Aop represents a potential node of crosstalk between the N and EGFR-MAPK signaling pathways. Though disputed, several reports indicate that EGFR signaling is required for the proper individuation of R8s from IGs (Baonza et al., 2001; Lesokhin et al., 1999; Lim and Choi, 2004; Chen and Chien, 1999). Early EGFR activation, through either mutant or transgenic means, elicits severe Ato defects (Lesokhin et al., 1999; Chen and Chien, 1999). Additionally, hypermorphic *Egfr* alleles are sensitive to reduced *aop* dosage, with such mutants featuring both Ato and eye defects (Rogge et al., 1995). In IGs, MAPK activation is dependent upon Ato and Da (Chen and Chien, 1999; Lim and Choi, 2004). This hints at a global mechanism of Ato regulation where repression of *ato* is primed but delayed due to the expression of a secondary factor, Aop. Aop is itself sensitive to EGFR-MAPK (Rebay and Rubin, 1995; Rogge et al., 1995). Thus, under this speculative mechanism, EGFR-MAPK would be required during R8 individuation to complement N signaling and permit simultaneous repression of both *ato* enhancers during Ato’s stage-3.

To clarify, Aop’s involvement in IG formation and R8 selection suggests that EGFR-MAPK is likely integrated into R8 selection through regulation of *ato-3’*. To maintain *ato-3’* activity during early stage-2, Aop must likely repress one or more genes that antagonize Ato. However, native Aop function would be impaired during stage-3 given by the activation of MAPK at that time. Thus, we expect Aop may regulate one or more genes that are 1) antagonistic to Ato expression, and 2) induced in or near the MF by MAPK signaling such as the homeorepressors Rough (Ro) and Bar (Dominguez et al., 1998; Lim and Choi, 2003).

The mechanism described herein bears homology to the signal-regulated events that coordinate human hematopoiesis. ETV6, the human ortholog of Aop, is expressed in multiple cell types of the hematopoietic stem cell lineage, as indicated by ETV6 association with myelodysplastic syndrome (MDS) (Bejar et al., 2011; Yoshika K, et al., 2011), acute myeloid leukemia (AML) (Barjesteh van Waalwijk van Doorn-Khosrovani et al., 2005), early T-cell acute lymphoblastic leukemia (ETP-ALL) (Van Vlierberghe et al., 2011a; Zhang et al., 2012), B-cell acute lymphoblastic leukemia (B-ALL) (Zhang J, et al., 2011), and diffuse large B-cell lymphoma (DLBCL) (Lohr et al., 2012). ETV6 was first implicated in leukemia for its dysregulation that was associated with transposition-derived fusion products; only recently has loss of ETV6 function become clearly linked to disease (Hock and Shimamura, 2018). Common lymphoid progenitors acquire T-cell fate through a combination of Notch and Ras signaling (Harman et al., 2003; Rothberg 2011; Lapinski and King, 2012). 15% of T-cell acute lymphoblastic leukemia (T-ALL) cases are comprised of a higher-risk form of the disease called ETP-ALL (Coustan-Smith et al., 2009). Of T-ALL subtypes, ETV6 mutations are vastly enriched in ETP-ALL (Van Vlierberghe et al., 2011b; Zhang et al., 2012). Although ETP-ALL, as a whole, features lower frequency of activational mutations to NOTCH1 when compared to general T-ALL, 80% of ETV6-mutant ETP-ALL cases are characterized by accompanying mutations in NOTCH1. Additionally, 67% of ETP-ALL cases had mutations in cytokine receptor and/or RAS signalling pathways, compared to only 19% of non-ETP cases (Zhang et al., 2011).

The role for ETV6 within T-cell specification is consistent with our proposed model for Aop’s function in R8 specification. In both cases, it is plausible that Notch signaling employs an ETS repressor to buffer the signaling environment from low levels of Ras, thus allowing Notch to set the minimum threshold of Ras activity that is required to trigger a response. In the event of R8 patterning, the Aop-maintained threshold ensures the imposition of a delay between the onset of Notch signaling and the onset of the repressive effects of Notch signaling, providing a critical window for the establishment of robust Ato autoregulation.

## Materials and Methods

### Drosophila Genetics

Flies were cultured on yeast-glucose media at 24°C and maintained according to a typical diurnal schedule. *aop^1^* is resultant of a nonsense mutation G952A that disables DNA binding (Caviglia and Luschnig, 2013). *aop^yan1^* (Bloomington Drosophila Stock Collection, BDSC, 8780) is a moderate loss-of-function allele that results from imprecise P-element excision from *aop^yanP^* that removes cytological map points 22D5-E1 (Lai and Rubin, 1992). *ato^1^* encodes A25T, K253N, N261I, the last of which ablates DNA binding (Jarman et al., 1994). *N^55e11^* (BDSC 28813) results from a 3.5kb insertion within the 5’ coding region of N, eliciting a premature nonsense mutation (Kidd et al., 1986). *Su(H)^^47^* results from an imprecise P-element excision that removes transcription start sites of Su(H) and l(2)Bg35 (Morel and Schweisguth, 2000). The rescue construct P{l(2)Bg35+} obviates contributions of l(2)Bg35 to Su(H) loss-of-function phenotypes described herein. *Df(3R)E(spl)^b32.2^* removes the entire *E(spl)* locus including *gro* (Schrons et al., 1992). The inclusion *of p{gro}* rescues cell-autonomous lethality caused by deletion of *gro* (Heitzler et al., 1996). *ato* reporter stocks, *5’-ato* (*P{w+mC ato5’F:9.3}* and *ato-3’* (*P{w+mC ato3’F:5.8}* are described in Sun et al., 1998. *h^H10^* results from a *pGawB* insertion in the *hairy* gene, which is used to drive *hairy*-dependent expression of GAL4 anterior to and within the MF (Ellis et al., 1994). UAS-mδ was made from the insertion of an EcoRI-XhoI fragment of *E(spl)mδ* cDNA into pUAST for forced-expression (de Celis et al., 1996; Ligoxygakis et al., 1999). dpp-lacZ is an enhancer reporter P-element insertion (Blackman et al., 1991; BDSC 5527 and 5528). dpp-GAL4 (BDSC 1553) was used in this study. UAS-aop^act^ (BDSC 5789) was first described in Rebay and Rubin, 1995.

### Immunohistochemistry

All steps were performed at room temperature unless otherwise indicated. Tissues were dissected in 0.1M sodium phosphate buffer and fixed in 4-6% formaldehyde, 0.1M sodium phosphate buffer. Tissues were washed in 0.3% Triton, 0.1M sodium phosphate buffer, 1% BSA, then blocked in 1% BSA, 0.1M sodium phosphate buffer, and incubated in primary antibody mixtures (antibody concentrations shown below in 0.1M sodium phosphate buffer) for 12-18 hours at 4°C. Following primary antibody incubation, tissues were washed in 0.1M sodium phosphate buffer and bathed in secondary antibody mixtures (1:1000 dilution for each secondary, in 1% BSA, 0.1M sodium phosphate buffer) for 2 hrs. Secondary antibody mixtures were removed, tissues were washed 0.1M sodium phosphate buffer. Tissues were mounted in 60% glycerol and imaged using an Olympus Fluoview FV1000 Confocal microscope. All scanning data reported was observed in a minimum of tissues from 5 independent animals of like genotype.

Primary antibodies include rabbit α-Ato (1:5000, (Jarman et al., 1994)); guinea pig α-Sens (1:500-800, (Nolo et al., 2000)); mouse α-E(spl)-mAb323 (1:3, (Jennings et al., 1994)); mouse α-β-gal-40-1a (1:800-1000); rat α-Ciact-2A1 (1:100, (Motzny and Holmgren, 1995)); mouse α-Aop (1:200, Rebay and Rubin, 1995); mouse α-Arm(1:100, Riggleman et al., 1990). mouse α-β-gal-40-1a, rat α-Ciact-2A1, mouse α-Aop, and mouse α-Arm were obtained from the Developmental Studies Hybridoma Bank, created by the NICHD of the NIH and maintained at The University of Iowa, Department of Biology, Iowa City, IA 52242. Tissues to be labeled with primary rabbit α-Ato were dissected in 0.3% Triton, 0.1M sodium phosphate buffer.

Secondary antibodies used include 488-goat α-mouse (Jackson), 488-goat α-Rat (Life Technologies), 546-goat α-rabbit (Life Technologies), 546-goat α-mouse (Life Technologies), 546-goat α-Rat (Life Technologies), 633-goat α-guinea pig (Life Technologies).

### Light Microscopy

Adult/pharate flies were mounted and promptly imaged using a Nikon camera in conjunction with a Leica MZ16 stereomicroscope, and eye size quantified as described (56). For counts listed as “700+”, facet count exceeded 700. Statistical significance was determined using Student’s T-Test.

### Image Production

All images were processed in Adobe Photoshop CC v. 14.2. Image manipulations of brightness/contrast and color balance were applied uniformly across each image shown. Images were then organized in Adobe Illustrator CC v. 17.1.

## ACKNOWLEDGMENTS

We are grateful to Drs. Nicholas Baker, Justin Kumar, Graeme Mardon, Yuh Nung Jan, Hugo Bellen, and Sarah Bray for generous gifts of flies and antibodies.

